# A molecular view into the structure and dynamics of phase-separated chromatin

**DOI:** 10.1101/2024.07.08.602582

**Authors:** Andrew Golembeski, Joshua Lequieu

**Affiliations:** Department of Chemical and Biological Engineering, Drexel University, Philadelphia, Pennsylvania 19104, USA

## Abstract

The organization of chromatin is critical for gene expression, yet the underlying mechanisms responsible for this organization remain unclear. Recent work has suggested that phase separation might play an important role in chromatin organization, yet the molecular forces that drive chromatin phase separation are poorly understood. In this work we interrogate a molecular model of chromatin to quantify the driving forces and thermodynamics of chromatin phase separation. By leveraging a multiscale approach, our molecular model is able to reproduce chromatin’s chemical and structural details at the level of nanometers, yet remain efficient enough to simulate chromatin phase separation across 100 nm length scales. We first demonstrate that our model can reproduce key experiments of phase separating nucleosomal arrays, and then apply our model to quantify the interactions that drive their formation into chromatin condensates with either liquid- or solid-like material properties. We next use our model to characterize the molecular structure within chromatin condensates and find that this structure is irregularly ordered and is inconsistent with existing 30 nm fiber models. Lastly we examine how post-translational modifications can modulate chromatin phase separation and how the acetylation of chromatin can lead to chromatin decompaction while still preserving phase separation. Taken together, our work provides a molecular view into the structure and dynamics of phase-separated chromatin and provides new insights into how phase separation might manifest in the nucleus of living cells.

## Introduction

Eukaryotic DNA must become highly compacted to fit within the confined space of the nucleus yet simultaneously organized to correctly facilitate gene expression. To meet these requirements, eukaryotic DNA complexes with histone octamers to form chromatin: a hierarchically compacted material which exhibits a heterogeneous and dynamic organization.^1^ Chromatin’s organization is driven by a diverse set of interactions between chromatin, RNAs, and proteins, resulting in several concurrent mechanisms which regulate chromatin’s organization.^2^

During interphase, chromatin is organized into distinct membraneless compartments like the nucleolus as well as heterochromatin and euchromatin domains. These compartments consist of different mixtures of DNA, RNAs, and proteins, and experiments have found that their constituents often interact to induce phase separation.^3^ Although the details of how phase separation might establish chromatin compartments is unclear,^4^ phase separation is now understood to play a role in the self-assembly and maintenance of membraneless bodies within the nucleus.^5^ For example, in the nucleolus liquid-liquid phase separation driven by weak, multivalent interactions is responsible for nucleolus formation as well as its management of nucleolar subcompartments.^6^ In heterochromatin, phase separation initiated by either HP1 bridging interactions or condensation explains how distal heterochromatic loci associate to form domains with regulated properties.^7-10^

Other recent work has focused on understanding the intrinsic phase separation of chromatin *in vitro*. In these studies, short chromatin segments consisting of reconstituted nucleosomal arrays have been observed to phase separate into dense chromatin condensates in the absence of chromatin-associated proteins and RNAs.^11-14^ These studies also provide clues to how chromatin phase separation may be regulated in the cell, such as the importance of linker histones and post-translational modifications in mediating chromatin phase separation.^11^

Nonetheless, many outstanding questions regarding the phase separation of chromatin remain. One open question relates to whether chromatin is liquid-like or solid-like and how this material state of chromatin might be modulated under different conditions. For example, recent experiments have observed that chromatin can be either liquid- or solid-like depending on the spacing of nucleosomes, the ionic conditions, and the presence of linker histones.^11-14^ Another open question relates to the mesoscale structure of chromatin condensates and whether there is any evidence for the solenoid and zigzag 30 nm fiber models.^15,16^ A final open question is how post-translational modifications, such as the acetylation of histone tails, can modulate the phase separation of chromatin across various length scales. Vhile recent simulation studies have made progress in starting to answer these questions,^17^ a detailed understanding of how these many factors can influence chromatin phase separation has not yet been established.

In this work, we investigate these questions by employing a molecular model of chromatin capable of simulating its phase separation directly at nucleosome resolution. Our results indicate that chromatin phase separation is driven by heterogeneous self-interactions which give rise to liquid-like behaviors on short length and timescales. However, we find that chromatin’s material state is sensitive to post-translational modifications and that solid-like chromatin can form when internucleosome interactions are increased. Our results also show that chromatin condensates lack a regular mesoscale structure like the solenoid and zigzag 30 nm fiber models, and that instead chromatin assumes a more disordered and irregular structure. Lastly we examine how H4 tail acetylation disrupts chromatin condensates and how H4 acetylation can decompact chromatin while still preserving it ability to phase separate. Taken together, our results provide a molecular view into chromatin condensates and how chromatin’s self-interactions can establish its organization.

## Methods

### Simulating chromatin condensates

We use the 1CPN model^18^ to simulate nucleosomal arrays and their phase separation. 1CPN is a coarse-grained, multiscale model that models linker DNA as a negatively charged twistable worm-like chain and nucleosomes as anisotropic sites with implicit histone tail interactions. These histone tail interactions are incorporated into 1CPN through coarse-grained potentials whose parameters are obtained using detailed free energy calculations from the near-atomistic 3SPN-AICG nucleosome model^19-24^ (see Ref. 18 and details below). This multiscale procedure permits the 1CPN model to be efficient enough to perform megabase-scale simulations of genomic DNA while retaining biochemical details such as post-translational modifications. 1CPN has been to shown to predict chromatin’s mesoscale structural features like tetranucleosomal folding motifs^25,26^ and is compatible with a recent model of the linker histone.^27^ Within 1CPN, electrostatics are treated using Debye-Hückel theory and the solvent is treated implicitly using Langevin dynamics.

Because 1CPN is parametrized using detailed 3SPN-AICG free energy calculations, the 1CPN model can be used to examine specific DNA sequences and post-translational modifications. In this work we use the strongly binding 601 DNA sequence and both unmodified nucleosomes and those with acetylated H4 tails. Specifically, the 601 DNA sequence is incorporated into 1CPN by matching its free energy profile for DNA’s rotation around the nucleosome to a profile calculated with 3SPN-AICG. Similarly, H4 tail acetylation is simulated by first computing anisotropic nucleosome pair potentials using the 3SPN-AICG model^24^ and then fitting 1CPN’s interaction parameters so that these potentials are reproduced.^18^ Additional details of this fitting process are provided in the supporting information.

We simulate the phase separation of nucleosomal arrays using the so-called direct coexistence method where both chromatin-dense and chromatin-dilute phases are allowed to equilibrate within a single simulation box. In order to mitigate the effects of the interface between phases, we adopt a slab-shaped simulation box which has been shown to reduce the finite-size effects present in direct coexistence simulations compared to cubic boxes.^28^ To initialize our system, we first place the nucleosomal arrays randomly in a large cubic simulation box and then compress the box until the x- and y-dimensions of the slab-shaped box are reached. The simulation box is then expanded along its z-dimension in order to form the slab-shaped box, and the system is then run until the dense and dilute phase concentrations equilibrate. We confirmed that these concentrations were not influenced by any of the specific parameters used in our initialization procedure (see supporting information). Uulk simulations of the chromatin-dense phase (i.e. chromatin condensate) were performed with cubic simulation boxes and a nucleosome concentration equal to that of the chromatin-dense phase.

### Classifying liquid- and solid-like chromatin condensates

Chromatin condensates are viscoelastic materials whose liquid- and solid-like properties depend on the length and timescales over which they are observed. In order to classify whether chromatin condensates are liquid- or solid-like we use two complementary metrics. For the first, we calculate the mean-squared displacement MSD) of nucleosomes as a function of time:^29^

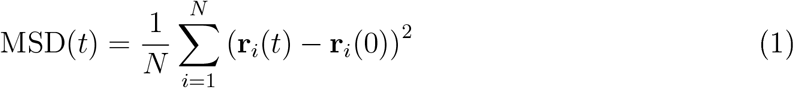

where **r**_*i*_(*t*) is the position vector of the *i*th nucleosome at time *t* and *N* is the total number of nucleosomes. Uy examining the slope of the MSD for a specified lag time, chromatin’s material state at different timescales can be obtained. For example, a MSD which scales linearly with time (indicated by a slope of 1 on a log-log plot) denotes freely diffusive nucleosomes characteristic of liquid-like chromatin. Alternatively, MSDs with slopes less than one denote subdiffusive nucleosome dynamics and more solid-like behavior.

The second metric we use to quantify the material state of chromatin examines the lifetime of nucleosome-nucleosome contacts within a condensate. To define this metric, we first specify that two nucleosomes with positions **r**_*i*_ and **r**_*j*_ are in contact if their spatial separation |**r**_*i*_ − **r**_*j*_| is less than the cutoff *r*_*c*_ = 150 Å. We then measure how these contacts are disrupted over time. Specifically, we define the fraction of persistent contacts *P* (*t*) as

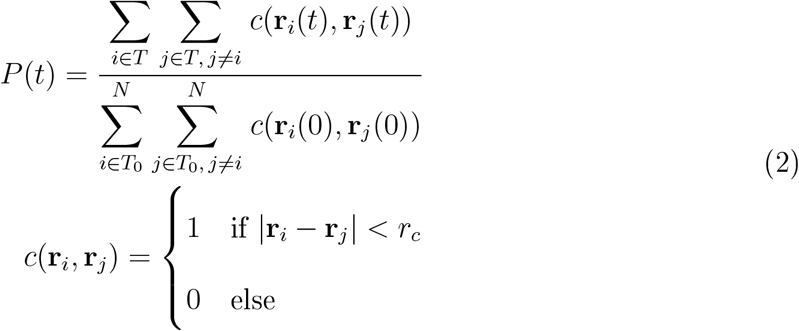

where *T*_0_ is the set of nucleosomes in contact with another nucleosome at time *t* = 0 and *T* is the set of nucleosomes which have persisted in their contact since *t* = 0. The timescale over which *P* (*t*) decays can be used to quantify the lifetimes of nucleosome-nucleosome contacts and corresponds to a transition from solid-like to liquid-like behavior. For both of these metrics, time is given in units of *τ*, the timescale over which a 12-mer nucleosomal array with a linker DNA length of 2,5 bp will diffuse its radius of gyration. This calculation and a further interpretation of *τ* is given in the supporting information.

### Characterizing structures within chromatin condensates

Chromatin structures such as the solenoid and zigzag 30 nm fiber models are defined by the interactions and orientations of their neighboring nucleosomes. To examine if solenoid and/or zigzag models are present within our chromatin condensates, we first define a pair of nucleosomes to be *k*th neighbors if they exist on the same chromatin chain and have *k* − 1 nucleosomes between them. We then define the orientation of two nucleosomes *i* and *j* using three angles: *α* which is the angle between the unit vectors normal to each nucleosome’s top face 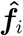 and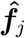, *β*_*i*_ which is the angle between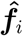 and the unit vector connecting the nucleosomes’ centers 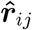, and *β*_*j*_ which is the angle between 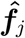 and 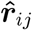.

Using these definitions, we can then classify nucleosome interactions by their different types: “face-face”, “face-side”, or “side-side” as described previously^17^ (see Table 1). “face-face” type nucleosome interactions between 1st and 2nd neighbors are characteristic of solenoid and zigzag 30 nm fiber models respectively,^30^ and their prevalence therefore quantifies the presence of these 30 nm fiber models within our simulations.

**Table 1:**
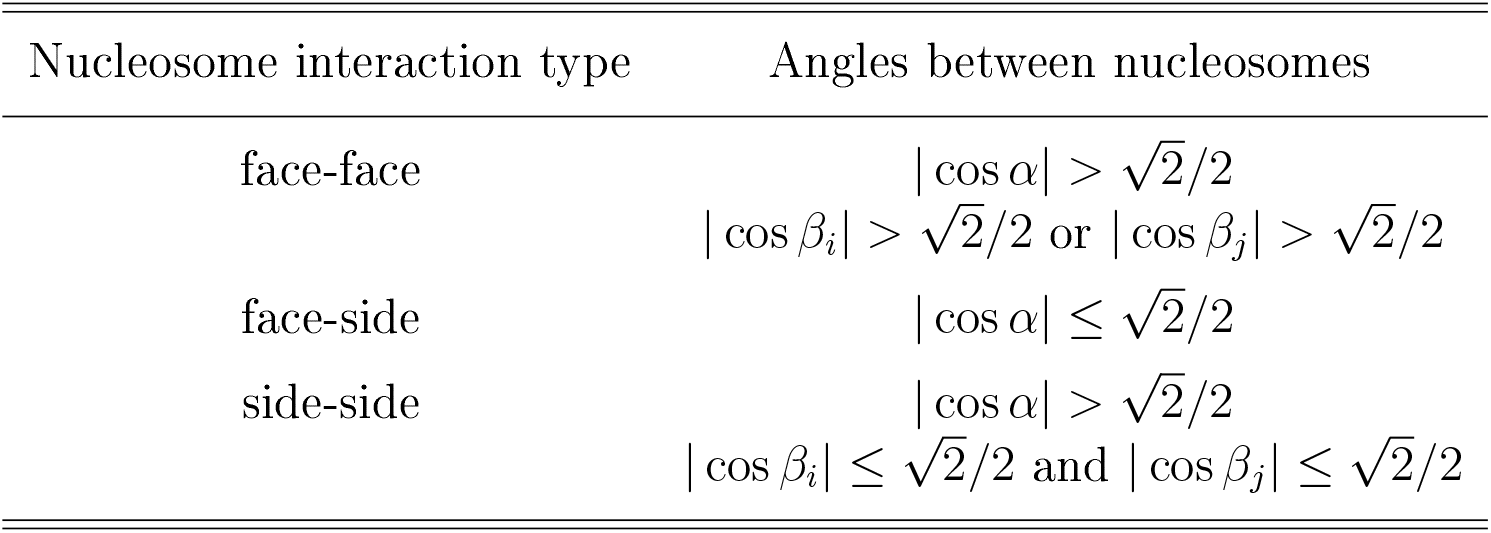
Definitions for nucleosome interaction types.

We also quantify the structure of nucleosomal arrays by measuring their diameters and linear packing densities. To do this, we approximate each array’s shape with a cylinder by using the following procedure. First, we calculate a nucleosomal array’s radius of gyration tensor *S*:^31^

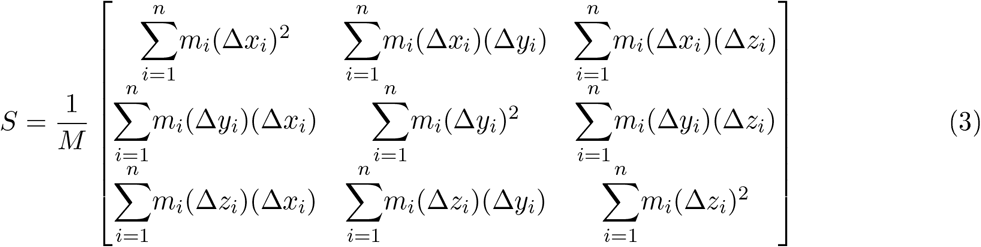

where *M* is the total mass of the array, *m*_*i*_ is the mass of an individual site *i, n* is the array’s number of sites, and Δ*x*_*i*_ is the distance between site *i* and the array’s center of mass along the *x*-axis, and similarly for the *y*- and *z*-axes. Once *S* is obtained, we then decompose *S* into eigenvectors *υ*_1_, *υ*_2_, and *υ*_3_ with corresponding eigenvalues *λ*_1_, *λ*_2_, and *λ*_3_ where *λ*_1_ ≥ *λ*_2_ ≥ *λ*_3_, and then define *υ*_1_ as the central axis of the approximating cylinder. Finally the cylinder’s length is then determined by the minimum and maximum positions of the array’s sites along its central axis, and the cylinder’s diameter is set equal to 2 times the maximum distance of the array’s sites from its central axis. When performing this procedure, we found that only using the single nucleosome site present in 1CPN underestimated the excluded volume of the nucleosome. This issue was easily rectified by introducing additional ghost sites which collectively represent the nucleosome’s excluded volume (see the supporting information).

The last metric we use to quantify chromatin’s structure within condensates is the nucleosome-nucleosome structure factor *S*(*k*), which is obtained by performing a Fourier transform on the nucleosome-nucleosome radial distribution function *g*(*r*) see Ref. 32):

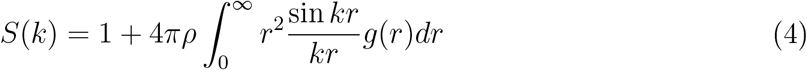

where *r* is a given radial distance, *k* is the magnitude of a given scattering vector, and *ρ* is the bulk nucleosome density. The magnitude of a scattering vector is related to a corresponding radial distance *r* = 2*π/k*, meaning that a structure factor peak at *k* = 2 indicates that nucleosome positions are correlated over distances of *r* = *π*.

## Results and discussion

Before interrogating the structure and thermodynamic properties of chromatin condensates, our first step is to validate our model and confirm that model predictions are consistent with existing experimental measurements. We validate our model using experimental data from Gibson et al. where dilute solutions of short nucleosomal arrays are observed to undergo phase separation. This seminal work has comprehensively mapped out the phase diagram of short nucleosomal arrays and has examined how phase boundaries can be modulated by numerous factors such as linker DNA length, salt concentration, the presence of linker histone H1, and histone acetylation. If our model can reproduce this wide range of experimental data, then we can have increased confidence in other model predictions for which experimental data is not available.

The first validation of our model focuses on experimental data from Gibson et al. which indicates that longer linker DNA lengths weaken chromatin phase separation, leading to lower concentrations of nucleosomes within the condensate. Gibson et al. also observe that the addition of linker histone H1 counteracts this effect and results in condensate nucleosome concentrations that are insensitive to linker DNA length. To examine if our model can reproduce these observations, we perform simulations that mirror the conditions used in the experiments: we simulate 12-mer nucleosomal arrays in solution with different linker lengths, in the presence and absence of linker histones, and examine whether these arrays undergo phase separation. If phase separation occurs, the nucleosome concentration within the condensate is measured. Our model predictions are in good agreement with experimental measurements (Fig. 1). Our model accurately predicts that the nucleosome concentration within condensates decreases for larger linker lengths in the absence of H1 but is constant if H1 is present.

**Figure 1.**
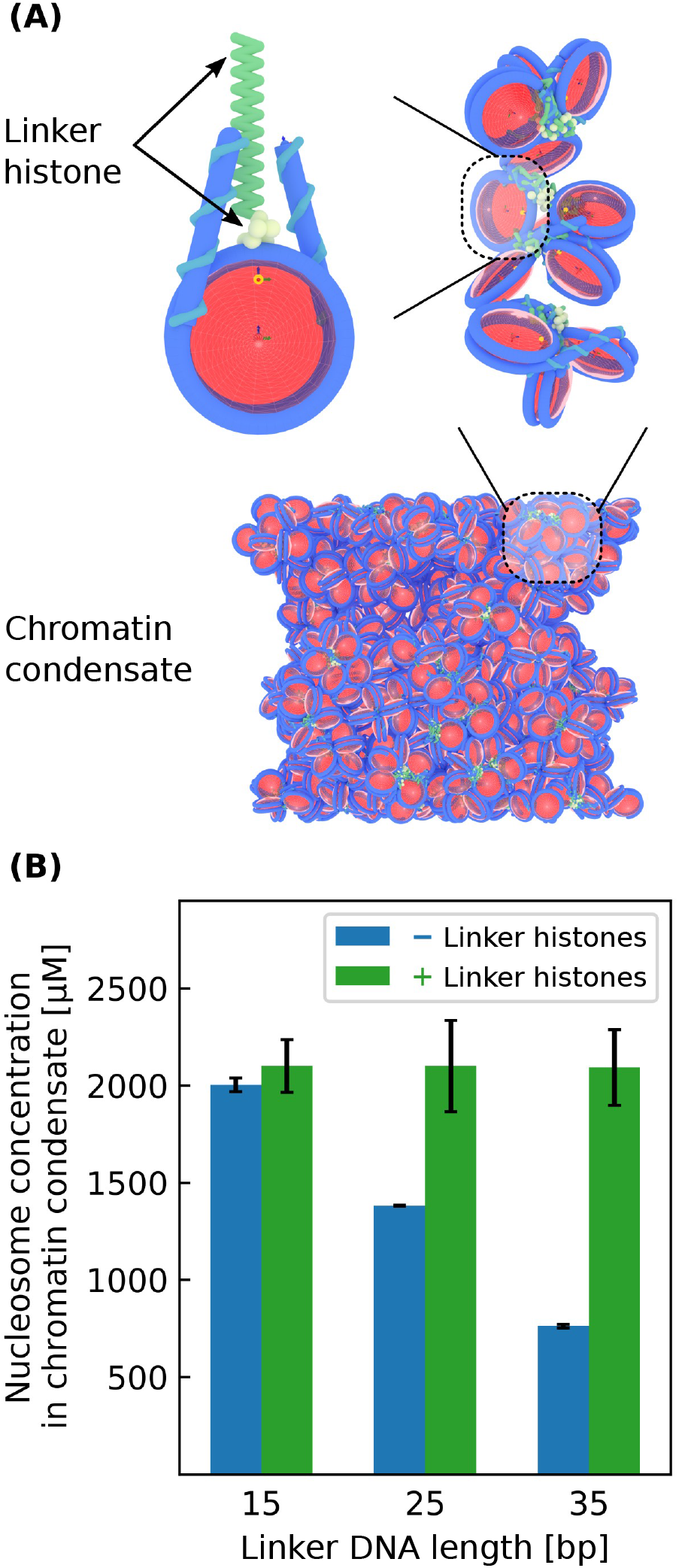
Phase separation of 12-mer nucleosomal arrays with varying linker DNA lengths with and without linker histones. **(A)** Model visualization of a chromatin condensate containing linker histones. **(B)** Nucleosome concentrations within the various chromatin condensates. Each simulation has a bulk nucleosome concentration of 200 µM and a box geometry of 100 x 100 x 1000 nm.

The next validation of our model focuses on other data from Gibson et al. where it was observed that lower salt concentrations and shorter nucleosomal arrays lengths i.e. having fewer nucleosomes per array) weaken chromatin phase separation. To test if our model can reproduce these results, we use our model to calculate the phase diagram for nucleosomal arrays with varying salt concentrations and nucleosomal array lengths. In these simulations the linker DNA lengths are fixed at 25 bp. Our model predictions agree well with experiments: while 12-mer solutions at high salt concentrations readily phase separate, solutions with salt concentrations less than 70 mM are incapable of phase separation (Fig. 2A). Similarly, while 12-mer nucleosomal arrays are capable of phase separation at a salt concentration of 150 mM, shorter 4-mer nucleosomal arrays are not (Fig. 2B).

**Figure 2.**
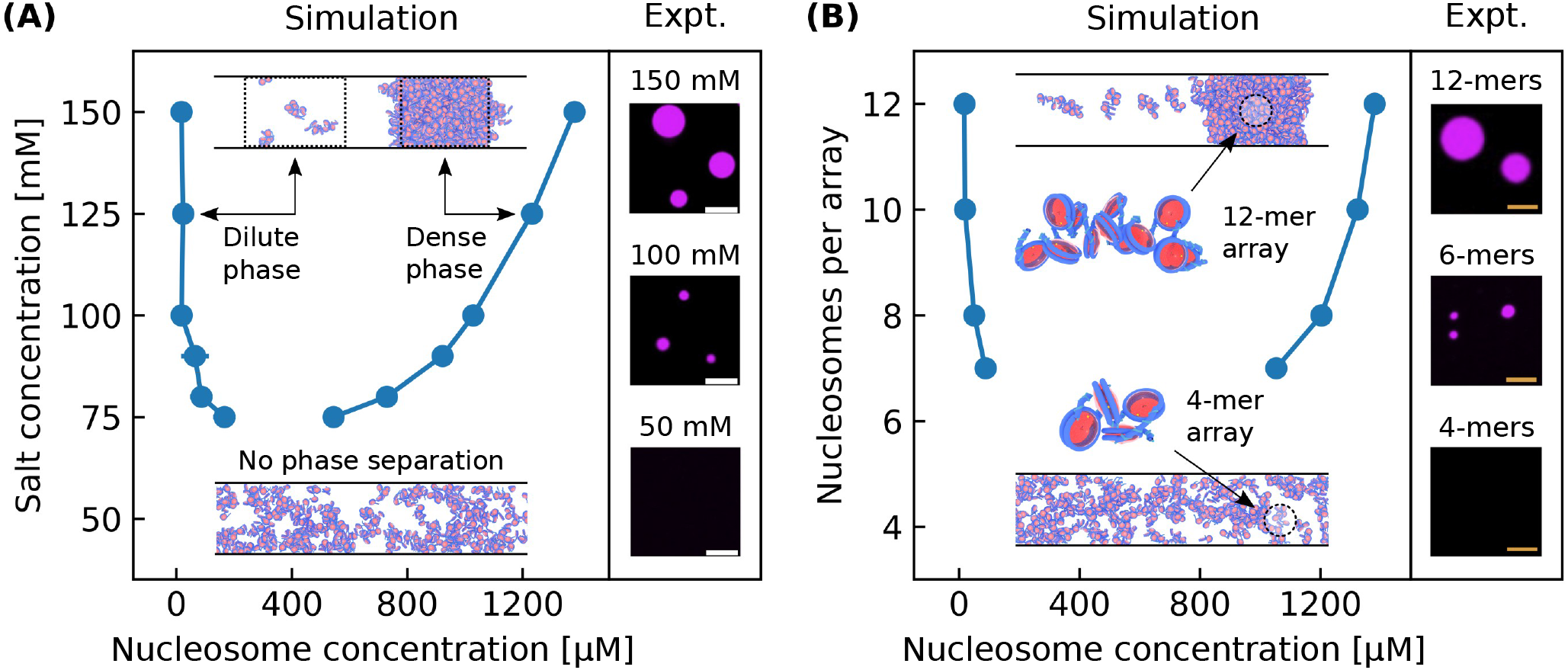
Phase separation of nucleosomal arrays under varying salt concentrations and nucleosomal arrays lengths. **(A)** Phase diagram with respect to salt concentration for 12-mer nucleosomal arrays alongside experimental results. **(B)** Phase diagram with respect to nucleosomal array length for nucleosomal arrays with a salt concentration of 150 mM alongside experimental results. Each simulation has a bulk nucleosome concentration of 200 µM and a box geometry of 100 x 100 x 1000 nm. Experimental data is from Ref. 11.

We emphasize that these simulation results are purely predictive and that no modifications were made to the originally published 1CPN model.^18^ As a consequence, the good agreement between simulation and experiment is quite remarkable, especially considering that the conditions within a chromatin condensate are very different from the dilute conditions under which 1CPN was originally parametrized.^18^ This parameter-free validation of our model gives us confidence that the 1CPN model can accurately predict the phase separation of chromatin condensates and that 1CPN can be used to provide molecular insights into their structural and material properties.

### Liquid- and solid-like chromatin condensates

Now that we have validated our model, we next turn to examine the conditions under which chromatin exhibits liquid- or solid-like material properties. Experiments have shown chromatin to be liquid-like^11,13,14^ or solid-like^12,33,34^ in various *in vitro* and *in vitro* systems, and these experiments collectively demonstrate that chromatin has a material state dependent on the length and timescales considered (i.e. chromatin is viscoelastic^35^). Chromatin’s material state is likely important to nuclear processes like transcription, but its significance and molecular origins are still poorly understood.

In order to examine how molecular interactions establish chromatin’s material properties, we focus on the role of internucleosome interactions. It is well-established that posttranslational modifications can modulate internucleosome interactions to make them weakly or strongly attractive.^36^ Wile we will examine the importance of specific post-translational modifications in a later subsection, we begin here by examining the role of internucleosome interactions at a coarse-grained level by varying the *ϵ*_0_ parameter in the 1CPN model (Fig. 3A), which scales the overall strength of the anisotropic interactions between nucleosomes see Eq., 5 in Ref. 18).

**Figure 3.**
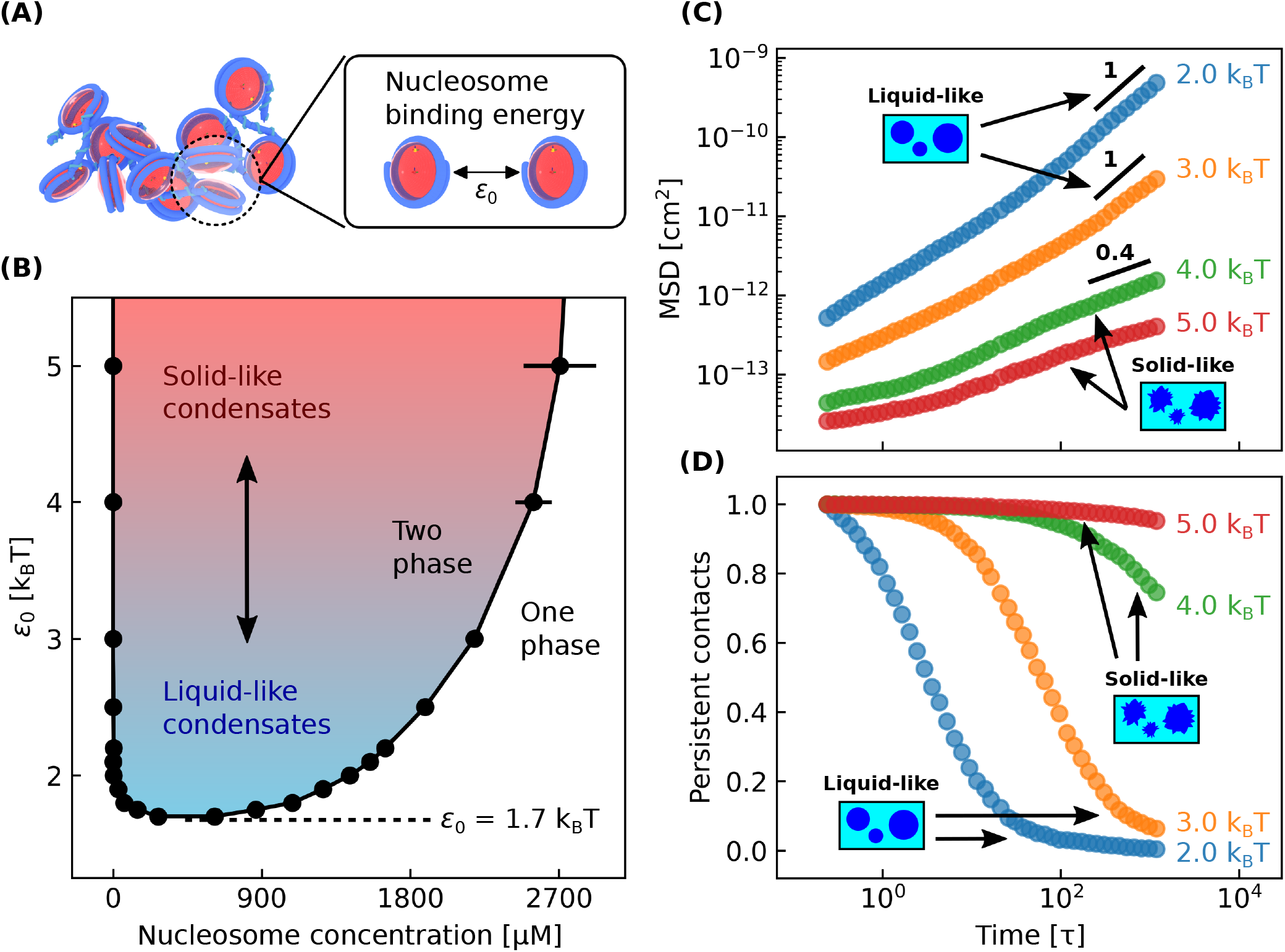
Liquid- and solid-like chromatin condensates. **(A)** Model visualization of the nucleosome binding energy *ϵ*_0_ which scales all internucleosome interactions. **(B)** Phase diagram with respect to *ϵ*_0_ for 12-mer nucleosomal arrays with linker DNA lengths of 25 bp. *ϵ*_0_ has a default value of 2.0 k_B_T in our model. **(C)** Mean-squared displacement MSD) over time for nucleosomes within different condensates. Liquid-like condensates have diffusive nucleosomes which linearly scale their MSD with time. **(D)** Fraction of persistent nucleosome contacts over time within different condensates. Vhen the persistent contacts equals 0, all initial nucleosome-nucleosome contacts are broken. *τ* is the time it takes a nucleosomal array simulated here to diffuse its radius of gyration and is equal to 40 µs. All simulations have a salt concentration of 150 mM. Simulations in **(B)** have bulk nucleosome concentrations of 300 µM and box geometries of 80 x 80 x 1000 nm. Simulations in **(C)** and **(D)** have bulk nucleosome concentrations equal to the corresponding chromatin-dense phase found in **(B)** and each contain 100 nucleosomal arrays.

To examine the effect of *ϵ*_0_ on chromatin’s material properties, we first compute the phase diagram for nucleosomal arrays as a function of *ϵ*_0_ (Fig. 3U). As anticipated, the phase diagram is very sensitive to *ϵ*_0_, with increasing values of *ϵ*_0_ resulting in a wider phase diagram and more strongly segregated chromatin. Notably, the phase diagram has a critical point of about 1.7 k_B_T which indicates that even relatively weak internucleosome interactions are sufficient to drive chromatin phase separation.

We next observe that the material state of chromatin is also very sensitive to *ϵ*_0_. We characterize chromatin’s material state within different condensates by computing the meansquared displacement (MSD) over time of their nucleosomes (Fig. 3C). Initially the MSD scales sublinearly with time which denotes subdiffusive nucleosomes typical of a solid-like chromatin state. As time progresses, the MSD eventually scales linearly with time which indicates liquid-like behavior. Over the timescale of our simulations (approximately 40 ms), we find that condensates with *ϵ*_0_ ≤ 3.0 k_B_T exhibit liquid-like behavior, while condensates with *ϵ*_0_ ≥ 4.0 k_B_T remain solid-like.

It is possible that diffusive nucleosomes have long-lived contacts characteristic of a solidlike material, and so we check for this by measuring the lifetime of nucleosome-nucleosome contacts within these condensates (Fig. 3D). We measure this lifetime by calculating the fraction of persistent contacts over time, which equals zero when all initial contacts are broken see Classifying liquid- and solid-like chromatin condensates). Fig. 3CD shows that the transition from subdiffusive to diffusive nucleosomes coincides with the termination of initial nucleosome-nucleosome contacts, and so we conclude that diffusive nucleosomes characterize a liquid-like chromatin state.

Overall we find that internucleosome interactions play a major role in establishing chromatin’s material properties. For example, by increasing internucleosome interactions by a mere 2 k_B_T, we observe that chromatin which was initially liquid-like will become solid-like (Fig. 3). Experimentally, the precise value of internucleosome interactions is the subject of some debate.^37-41^ Some recent experiments have estimated internucleosomal attractions to be at most 2.7 k_B_T,^41^ which based on our results suggests that short segments of unmodified chromatin are liquid-like in agreement with some recent experiments^11,14,42^ but not others.^12^ Since chromatin’s material state is sensitive to *ϵ*_0_, it is likely that changes to *ϵ*_0_ through post-translational modifications can modulate the timescales over which chromatin is liquid- or solid-like. Such regulation is likely important for transcription which occurs over similar timescales of 10-100 ms per nucleotide.^43^

### Mesoscale structure of chromatin condensates

We now turn to characterize chromatin’s mesoscale structure within chromatin condensates. Our first objective is to investigate whether solenoid or zigzag 30 nm fibers^44^ are present in our simulations. These structures can be identified by the way their neighboring nucleosomes interact^30^ and so we begin by analyzing these interactions. To carry out this analysis, we first perform a bulk simulation of a liquid-like chromatin condensate (Fig. 4A) and then measure the frequency of different interaction types between 1st and 2nd nucle-osome neighbors (Fig. 4U). Our results indicate that liquid-like chromatin contains mostly face-side interactions between 1st and 2nd neighbors and that face-face interactions rarely occur. Since both solenoid and zigzag models are characterized by sustained face-face interactions, (Fig. 4U shows that neither solenoid nor zigzag fibers are present within liquid-like chromatin condensates.

**Figure 4.**
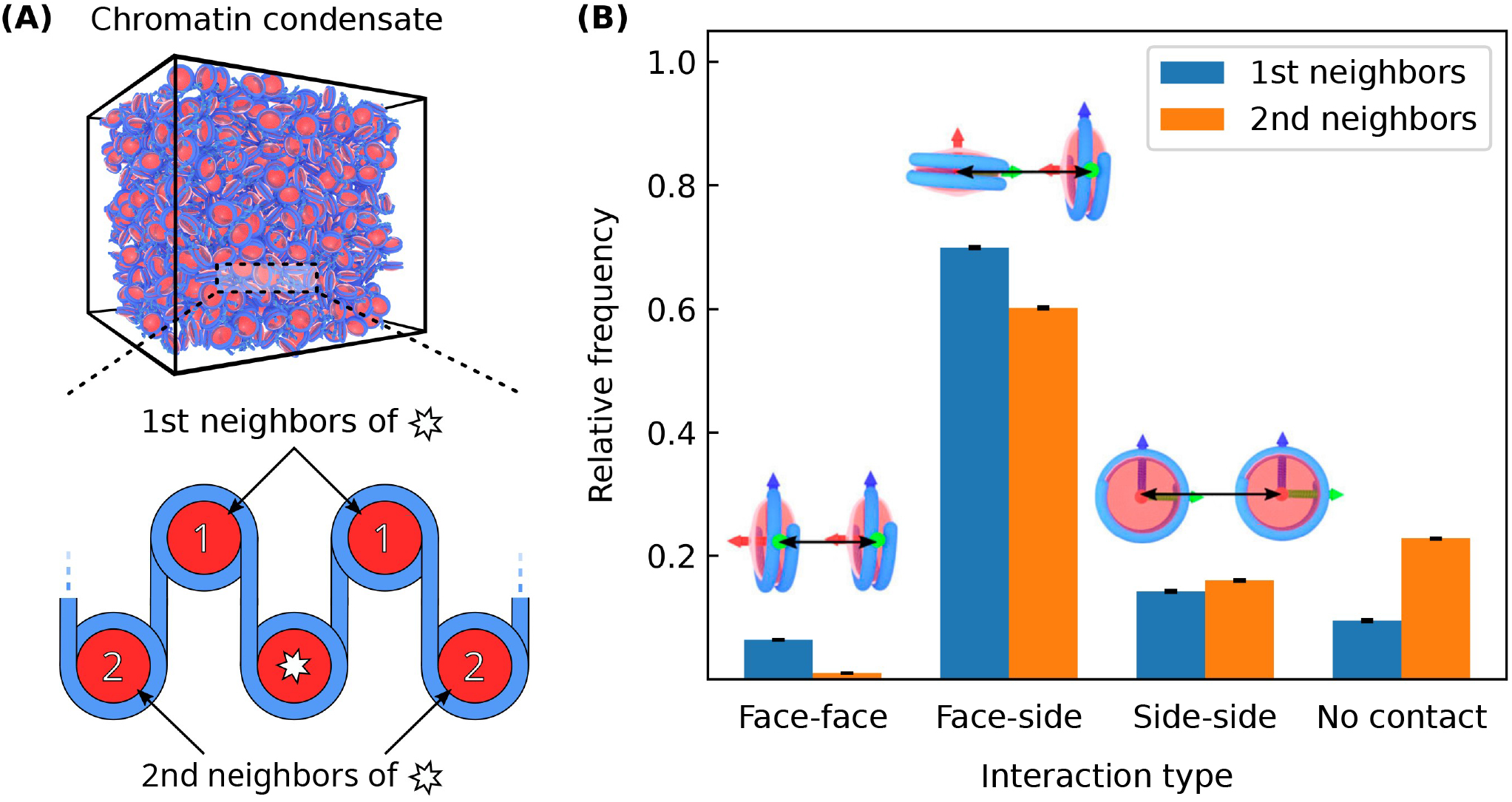
Interactions between neighboring nucleosomes for liquid-like chromatin (*ϵ*_0_ - 2.0 k_B_T). **(A)** Model visualization of a bulk chromatin condensate simulation top) and a cartoon depicting nucleosome neighbors (bottom). **(B)** Relative frequencies of different interaction types between neighboring nucleosomes. Visualizations of each interaction type are shown above their respective values. Neighboring nucleosomes more than 150Å apart are not interacting and have interaction type “no contact”. The bulk condensate simulation contains 100 12-mer nucleosomal arrays having linker DNA lengths of 25 bp with a salt concentration of 150 mM and bulk nucleosome concentration of 1400 µM.

We next characterize chromatin’s mesoscale structure directly by fitting each nucleosomal array to a cylinder of minimal volume see Characterizing structures within chromatin condensates). This fitting allows us to quantify the diameter of each array and the corresponding distribution of array diameters throughout a chromatin condensate (Fig., 5A). Surprisingly, we observe that the average array diameter is around 30 nm despite the previous lack of evidence for 30 nm fiber models. However, the distribution of array diameters is quite wide which indicates that chromatin’s mesoscale structure is irregular throughout the liquid-like condensate. To better understand this broad distribution, we examine the linear packing densities of nucleosomal arrays which measures how tightly compacted each array is (Fig., 5U). The arrays’ packing densities are similarly widely distributed and the average packing density is roughly 2 times lower than that of either 30 nm fiber model. Together these distributions show that liquid-like chromatin has an irregular mesoscale organization different from the 30 nm fiber models. These results suggest that chromatin’s mesoscale structure is instead better described as a disordered polymer chain which is qualitatively consistent with chromatin electron microscopy tomography (ChromEMT) experiments.^45^

**Figure 5:**
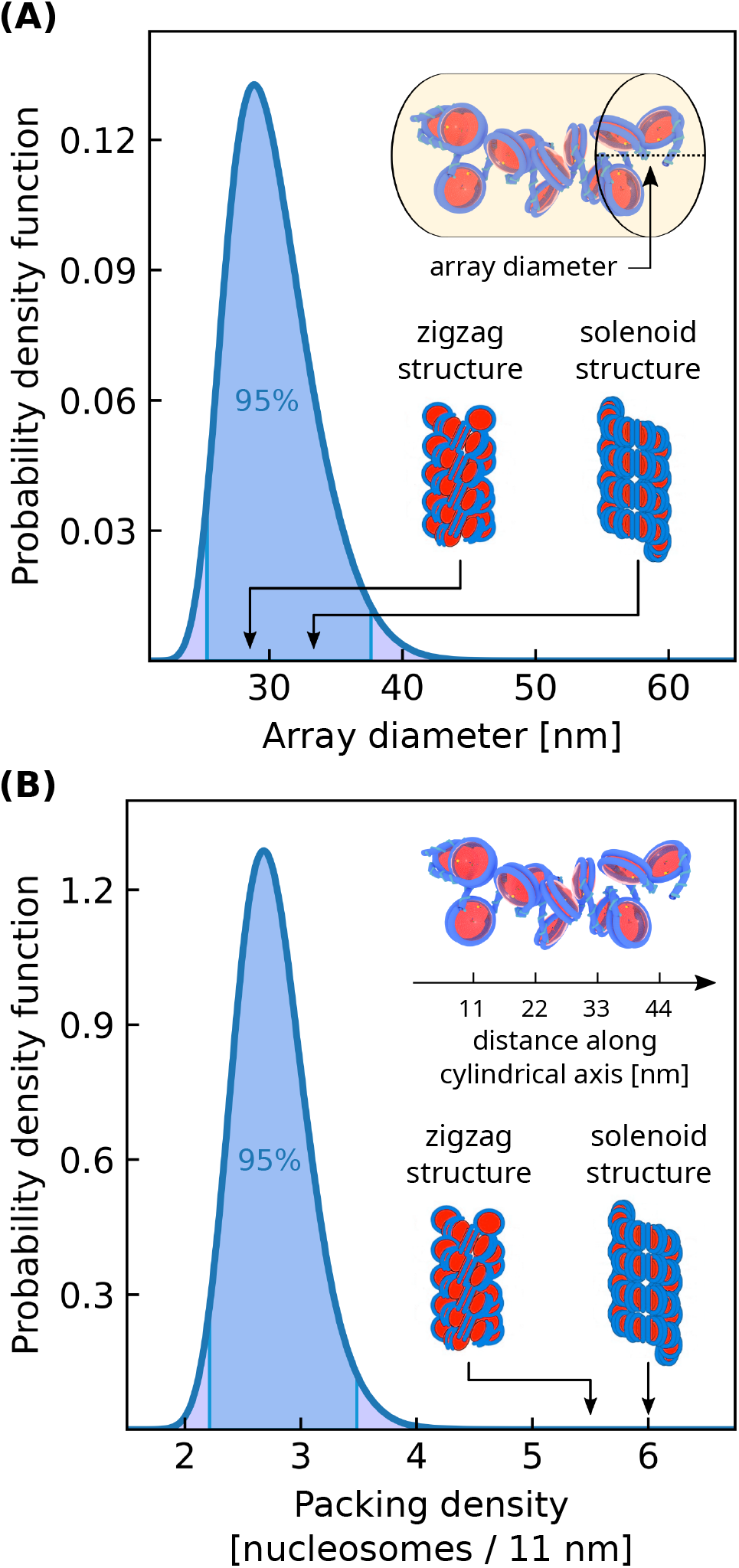
Nucleosomal array structures within a liquid-like chromatin condensate (*ϵ*_0_ = 2.0 k_B_T). (**A**) Distribution of array diameters. (**B**) Distribution of array packing densities. The packing density is defined as the number of nucleosomes per 11 nm along the array’s central axis. Values for solenoid and zigzag 30 nm fiber models are indicated using illustrations from Ref. 46. The bulk condensate simulation contains 100 12-mer nucleosomal arrays having linker DNA lengths of 25 bp with a salt concentration of 150 mM and bulk nucleosome concentration of 1400 µM.

Finally, we examine how the mesoscale structure of chromatin changes as a condensate varies from liquid-to solid-like. To perform this analysis, we first conduct bulk simulations of chromatin condensates with different internucleosome interaction strengths scaled by *ϵ*_0_, which we previously found to change chromatin’s material state (Fig. 3). We then calculate the nucleosome-nucleosome structure factor of each condensate (Fig. 6). For liquid-like condensates (*ϵ*_0_ = 2.0 k_B_T), the structure factor is peaked at 8.5 nm and shows no peak at 30 nm. The peak at 8.,5 nm corresponds to frequent face-side nucleosome interactions which occur over distances of 7-11 nm and is consistent with our observations in Fig. 4U. The absence of a peak at 30 nm is also consistent with the broad distribution of array diameters that we observed in Fig., 5A. As *ϵ*_0_ is increased and chromatin becomes more solid-like, the peak in the structure factor shifts to smaller distances and is peaked at 7.1 nm for *ϵ*_0_ = 5.0 k_B_T. This indicates that stronger internucleosome interactions result in more compact chromatin structures, consistent with Fig. 3U. Even at larger values of *ϵ*_0_, we see no peak at 30 nm which suggests that no well-ordered 30 nm fibers are present within our simulations. We also observe that larger values of *ϵ*_0_ decrease chromatin’s isothermal compressibility which is equal to the structure factor *S*(*k*) as *k* → 0.^47^

**Figure 6.**
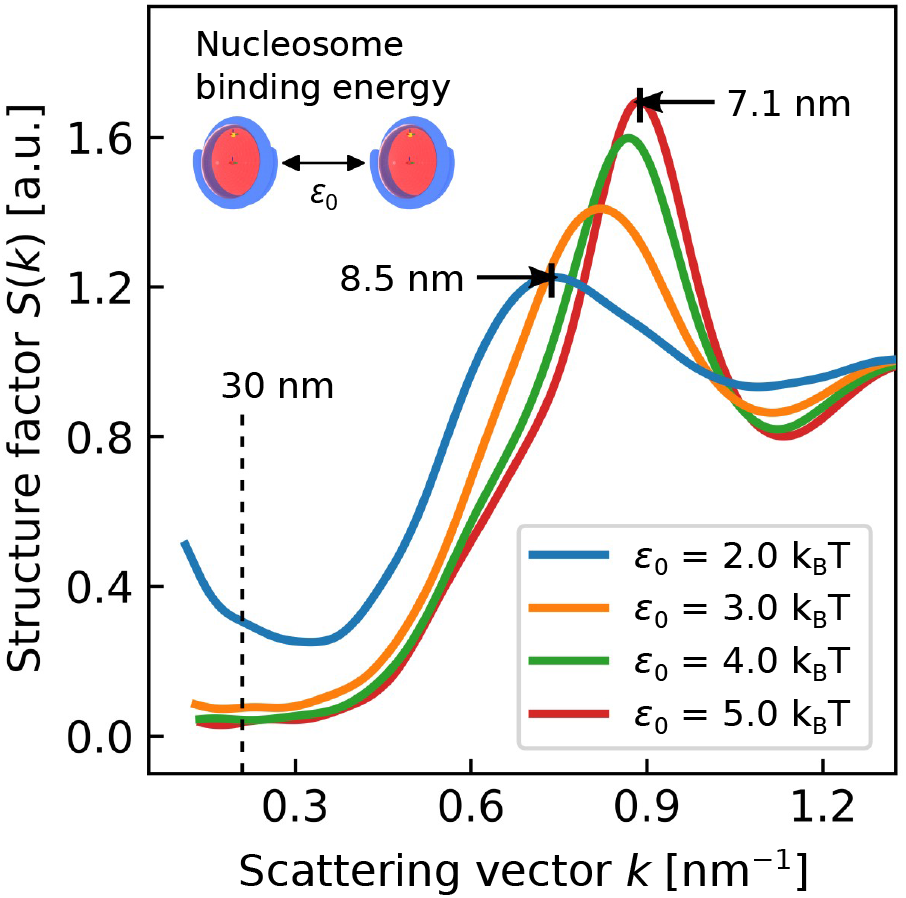
Nucleosome-nucleosome structure factor for liquid- and solid-like chromatin condensates. Structure factor peaks indicate correlated nucleosome positions over the given reciprocal scattering vector length 2*π/k*. Nucleosome binding energies *ϵ*_0_ ≤ 3.0 k_B_T correspond to liquid-like condensates while *ϵ*_0_ ≥ 4.0 k_B_T correspond to solid-like condensates. Each bulk condensate simulation contains 100 12-mer nucleosomal arrays having linker DNA lengths of 25 bp with a salt concentration of 150 mM and bulk nucleosome concentration equal to that of the condensate with nucleosome binding energy *ϵ*_0_.

By further increasing internucleosome interactions, our results suggest that the face-face nucleosome interactions of 30 nm fiber models will eventually become favorable. Such highly attractive internucleosome interactions have been explored in experiments of nucleosomal arrays with superphysiological concentrations of divalent salts.^14,30^ In agreement with our model predictions, these nucleosomal arrays were shown to phase separate into solid-like condensates^14^ and have frequent face-face interactions.^30^ Interestingly, these arrays did not regularly form into 30 nm fiber models despite their frequent face-face interactions.^30^ This may be explained by the 60 bp linker DNA lengths of the arrays which are much longer than the 25 bp linker lengths considered here. Longer linker DNA lengths drive chromatin’s structure to be more irregular and flexible^25^ and so altogether it is likely that the formation of 30 nm fibers is sensitive to both the nuclear environment and chromatin’s composition. This sensitivity can explain experimental results showing that 30 nm fibers are rare occurrences in human chromatin^48^ and common in chicken erythrocytes.^49^

### H4 acetylated chromatin condensates

We now turn to examine how specific post-translational modifications modulate chromatin phase separation. We focus here on histone acetylation which has been shown to weaken chromatin’s self-interactions^24,41^ and inhibit the phase separation of 12-mer nucleosomal arrays.^11^ Since chromatin’s H4 tails are essential for its compaction,^50^ we focus on how the acetylation of H4 tails (here referred to as H4 acetylation) can modulate chromatin phase separation.

In order to simulate H4 acetylated chromatin, we first compute the orientation-dependent pair potential between two H4 acetylated nucleosomes using the near-atomistic 3SPN-AICG model (Fig. 7A). This orientation-dependent pair potential is then used to reparametrize the 1CPN model so that it can incorporate the subtle effects of H4 acetylation (see Fig. 7UC and Simulating chromatin condensates). We note that this multiscale approach to incorporate post-translational modifications is essentially parameter-free and provides a systematic route to include chemically specific information into the 1CPN model.

**Figure 7.**
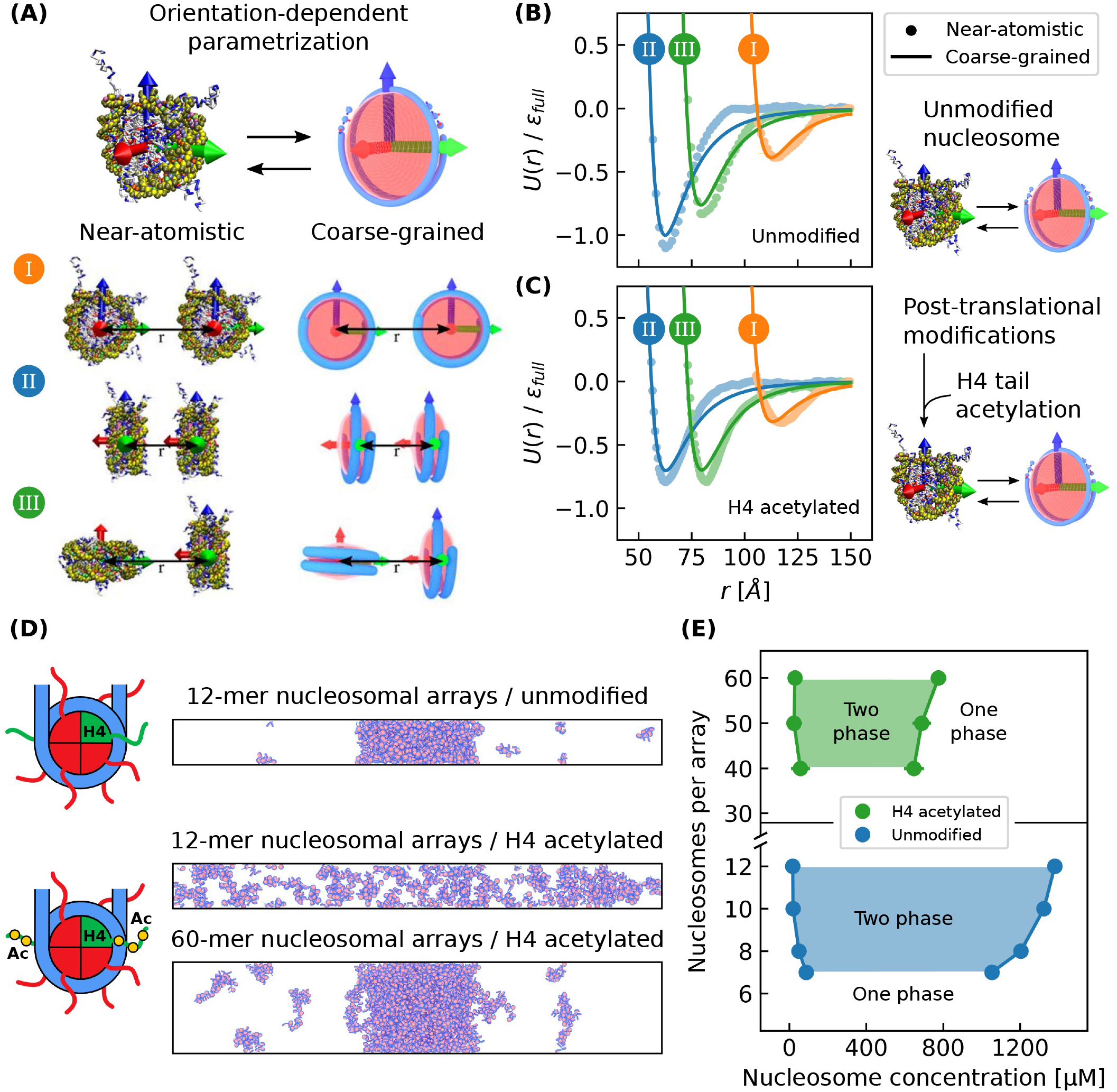
H4 tail acetylation of chromatin condensates. **(A)** The near-atomistic 3SPN-AICG nucleosome model was used to obtain orientation-dependent pair potentials which parameterize the coarse-grained 1CPN model. The orientations shown are (I) side-side, (II) face-face, and (III) face-side. **(B)** Near-atomistic and coarse-grained 1CPN nucleosome pair potentials for unmodified chromatin and **(C)** H4 acetylated chromatin. **(D)** Simulations of unmodified 12-mer nucleosomal arrays and H4 acetylated 12-mer/60-mer arrays with linker DNA lengths of 2,5 bp. **(E)** Phase diagram with respect to nucleosomal array length for unmodified and H4 acetylated chromatin.

We first apply our model of H4 acetylation to 12-mer nucleosomal arrays. Recent experiments have observed that this system should undergo phase separation for unmodified chromatin but should not when chromatin’s H4 tails are partially acetylated.^11^ Our model predictions are in agreement with these experiments and accurately predict that H4 acetylation is sufficient to disrupt the phase separation of 12-mer arrays (Fig. 7D). Our model also predicts that the phase separation of H4 acetylated chromatin can be restored by increasing its length to 60-mer arrays (Fig. 7D). The reemergence of phase separation for longer arrays can be rationalized by the longer arrays’ lower entropy of mixing which favors phase separation.^51^ The ability of longer stretches of acetylated chromatin to reinitiate phase separation suggests that chromatin phase separation in living cells is likely a multifaceted process where both the type and density of post-translational modifications can be used to tune phase separation.

To quantify the effect of H4 acetylation on chromatin phase separation, we compute a phase diagram with respect to array length for H4 acetylated chromatin and compare it to unmodified chromatin (Fig. 7E). We find that H4 acetylation shifts the phase diagram to larger array lengths, and that H4 acetylated chromatin requires 40-mer arrays to phase separate while unmodified chromatin only requires 7-mer arrays. We also find that H4 acetylation results in a lower concentration of nucleosomes in the chromatin-dense phase (650 - 850 µM) which is approximately half the concentration observed in unmodified chromatin (1050 - 1400 µM). Taken together, our results indicate that even though H4 acetylation can decompact chromatin,^42^ this decompaction does not exclude the possibility that acetylated chromatin can phase separate, especially for long chromatin chains. This result helps explain recent evidence suggesting that acetylated chromatin regions are liquid-like, phase separated domains^34^ and further supports the theory that phase separation can play a major role in chromatin’s organization.

## Conclusions

In this work we examine chromatin condensates composed of nucleosomal arrays by utilizing a multiscale, coarse-grained chromatin model. Our model can reproduce key experimental observations of these condensates including their dependence on linker histones, salt concentration, linker DNA length, and array length. Using this model, we have demonstrated that the material state of chromatin condensates can be modulated by the strength of nucleosome-nucleosome interactions. Based on experimental estimates of chromatin’s nucleosome-nucleosome interactions,^41^ our model predicts that unmodified chromatin is liquid-like on short length scales in agreement with experiments.^11,13,14^ Like other viscoelastic materials, chromatin transitions from liquid-to solid-like behavior depending on the length and timescale considered^34,35^ and our results demonstrate that post-translational modifications can regulate this transition for chromatin.

Our work also provides a molecular view into chromatin’s mesoscale structure which regulates gene expression yet is poorly defined.^16^ Within chromatin condensates, our results show that chromatin lacks a regular secondary structure like the solenoid and zigzag 30 nm fiber models. Instead chromatin is structured irregularly and frequently self-interacts through heterogeneous nucleosome-nucleosome interactions as also observed experimentally.^45^ We additionally show that chromatin’s mesoscale structure becomes more regular as chromatin’s self-interactions are increased, which can occur under superphysiological concentrations of divalent cations^30^ or when nucleosomes are more closely spaced.^25^

Lastly our work investigates whether H4 tail acetylation can prevent chromatin phase separation. In agreement with experiments,^11^ we find that H4 acetylation dissolves chromatin condensates composed of short, 12-mer nucleosomal arrays. However, we also observe that condensates of acetylated 40-mer nucleosomal arrays can reinitiate phase separation, albeit at a lower degree of compaction relative to unmodified chromatin. This result reveals that acetylation’s decompaction of chromatin is compatible with chromatin phase separation and also exemplifies how phase separation may play a major role in chromatin’s organization. Collectively, our work demonstrates that multiscale, coarse-grained models are a powerful tool for probing the structural and dynamic properties of *in vitro* chromatin systems.

## Acknowledgenent

A.G. was supported by the Department of Education under Award No. P200A180026. This work was also partially supported by a New Investigator Grant from the Charles E. Kaufman Foundation. Simulations were run on hardware supported by Drexel’s University Research Computing Facility.

## Supporting Information Available

The Supporting Information is available free of charge at URL.

