## Supporting information for "A molecular view into the structure and dynamics of phase-separated chromatin"

### Simulating chromatin phase separation

Each simulation was run using the 1CPN model which has an accompanying quick start guide located here. Unless otherwise stated, the default 1CPN parameters[S1] are used in our simulations. We simulate phase separation directly by using a slab-shaped simulation box which minimizes the effects of a finite-sized interface between phases.[S2] A "slab" here refers to a rectangular prism having two dimensions of equal length and one dimension of a greater length, which in our work is the $z$-axis. We prepare systems with slab-shaped geometries by first randomly initializing them in a large simulation box, in order to minimize any initial overlap of sites. We then compress the box into an intermediate slab-shaped geometry. This intermediate geometry is the same as the final geometry, except that the box's length along the $z$-axis is lower than its final value, and at this point a chromatin-dense phase (i.e. chromatin condensate) is formed. After the system equilibrates in this intermediate geometry, the box's length along the $z$-axis is then extended to its final value which provides volume for the chromatin-dilute phase. Afterwards 10 nucleosomal arrays are optionally displaced from the dense phase into the dilute phase in order to reduce equilibration time. We then run the simulation until the nucleosome concentration of each phase has reached a steady state value and at that time we consider the system equilibrated.

In order to calculate the nucleosome concentration of each phase, we construct a concentration profile along the box's $z$-axis which we use in the main text to construct binodals of chromatin phase separation. For example, the concentration profiles used to construct the binodal for unmodified 12-mer nucleosomal arrays with respect to the nucleosome binding energy $\epsilon_0$ are shown here (Fig. S1). From these profiles, we calculate the chromatin-dense and dilute phase concentrations by averaging the nucleosome concentration over their location along the $z$-axis.

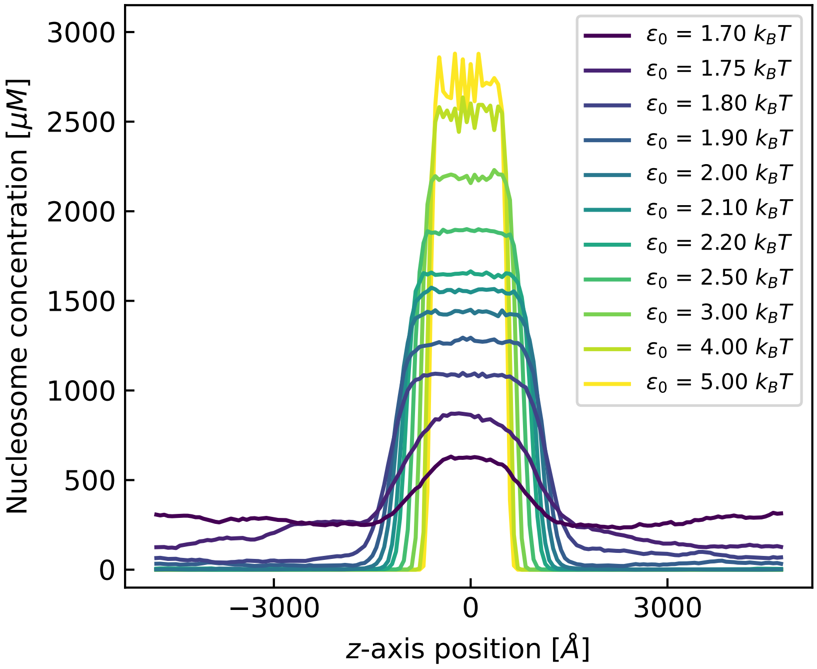


Figure S1: Nucleosome concentration profiles for direct coexistence simulations of 12-mer nucleosomal arrays with different nucleosome binding energies $\epsilon_0$. All simulations have a salt concentration of 150 mM, a bulk nucleosome concentration of 300 µM, and a box geometry of 800 x 800 x 10000 Å. Each nucleosomal array has a linker DNA length of 25 bp.

Because phase equilibria is simulated directly, the nucleosome concentration of each phase can be biased by the simulation box's dimensions, as it influences the simulated interface between the coexisting phases. The nucleosome concentration can also be biased by self-interactions across the box's periodic boundaries, which are dependent on the length of the nucleosomal arrays and the box's geometry. These biases are known as finite-size effects, and we find these effects to be minimal for slab-shaped box geometries larger than 792x792x9584 angstroms ($x$-axis x $y$-axis x $z$-axis) for systems containing 12-mer nucleosomal arrays with a linker DNA length of 25 bp (Fig. S2). Throughout our work, we seek to minimize the box's cross-sectional area ($x$-axis x $y$-axis) in order to reduce equilibration time while keeping it large enough to prevent self-interactions across periodic boundaries.

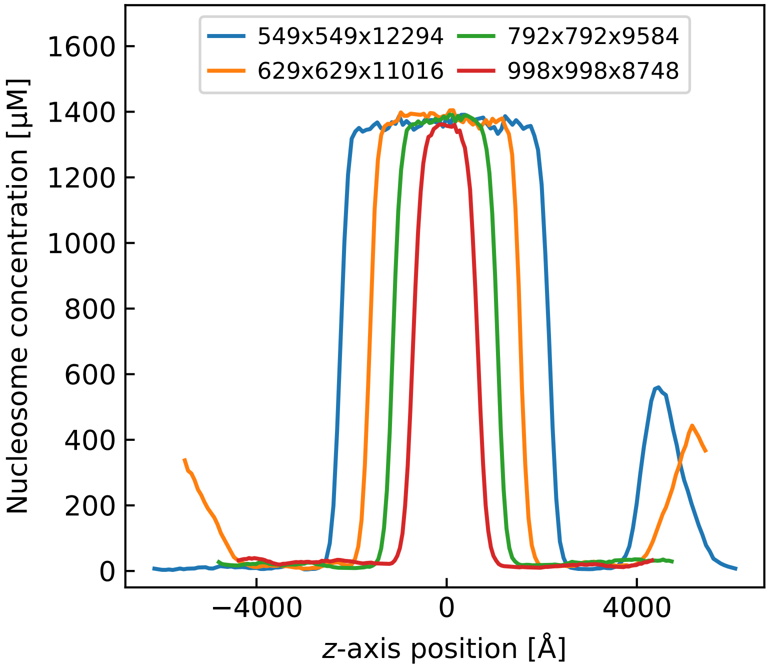


Figure S2: Nucleosome concentration profiles for direct coexistence simulations with different box geometries ($x$-axis x $y$-axis x $z$-axis). All simulations have a salt concentration of 150 mM and contain 100 12-mer nucleosomal arrays with linker DNA lengths of 25 bp.

Once the nucleosome concentration of a chromatin condensate is determined, we simulate the condensate in the absence of its corresponding dilute phase in order to measure its bulk properties. These so-called "bulk condensate simulations" allow us to more easily calculate the properties of the condensate while simultaneously minimizing interfacial biases. We prepare bulk condensate simulations simply by compressing a large, randomly initialized simulation box into a cubic box having a total bulk nucleosome concentration equal to that of the specified condensate. We then use the trajectories of these simulations to calculate properties like the mean-squared displacement of nucleosomes, the fraction of persistent contacts, the interaction types of neighboring nucleosomes, the diameters and packing densities of nucleosomal arrays, and the nucleosome-nucleosome structure factor.

### Simulating under a Langevin thermostat

1CPN samples the NVT ensemble by using a Langevin thermostat which implements freely draining Brownian dynamics that maintain the system's temperature and implicitly model solvent effects (see Ref. S1). A Langevin thermostat is parametrized by a damping parameter $\lambda$ which determines the viscosity of the implicit solvent and consequently the ratio of frictional forces to thermal fluctuations felt by each of 1CPN's sites. Because nucleosomes have a large excluded volume, the damping parameter for nucleosome sites is $6\lambda$ while for DNA and dyad sites it is $\lambda$. For higher values of $\lambda$, 1CPN's implicit solvent becomes less viscous which causes the simulation to more rapidly equilibrate, but for values of $\lambda$ too large the increased thermal fluctuations bias the simulation's equilibrium properties.

We explored this possible bias by calculating the nucleosome concentration profiles for systems with different values of $\lambda$ (Fig. S3). We found that $\lambda$ does not affect equilibrium properties like nucleosome concentrations within phases unless extreme values are chosen. We therefore set $\lambda = 5 \times 10^6$ for our simulations in order to have efficient simulations without introducing an artificial bias.

Importantly, because $\lambda$ controls the speed of the system's dynamics, Langevin thermostats cause absolute simulation timescales to be less meaningful than relative ones. In order to extract meaningful dynamic properties from our simulations, we nondimensionalize time with respect to a characteristic timescale $\tau$ which is the simulation time it takes for a 12-mer nucleosomal array with linker DNA length 25 bp to diffuse its radius of gyration in dilute conditions.

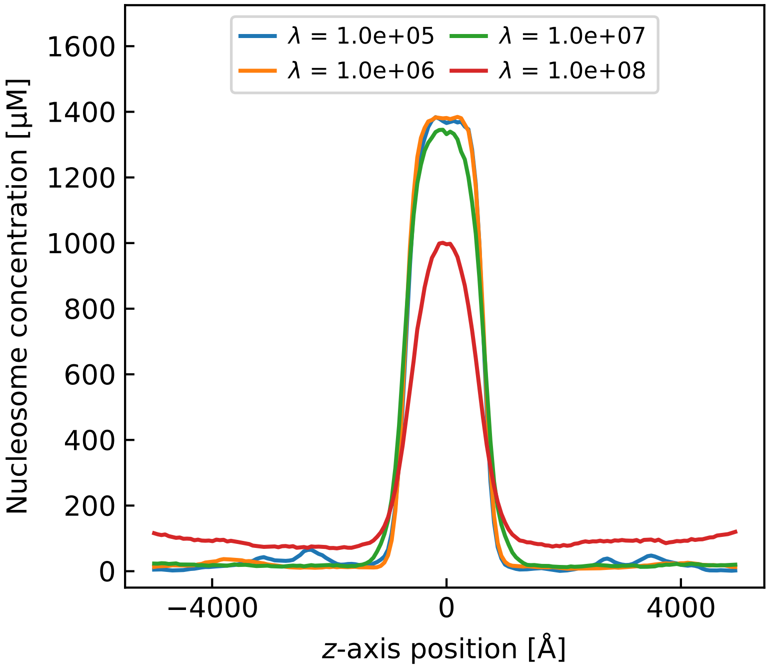


Figure S3: Nucleosome concentration profiles for direct coexistence simulations of 12-mer nucleosomal arrays with different Langevin damping parameters $\lambda$. All simulations have a salt concentration of 150 mM, a bulk nucleosome concentration of 200 µM, and a box geometry of 1000 x 1000 x 10000 Å. Each nucleosomal array has a linker DNA length of 25 bp.

To calculate $\tau$, we first simulate the 12-mer nucleosomal array in a large box and compute its radius of gyration $R_g$ averaged over each simulation frame, which we find to be $R_g =$ 145 Å. We then calculate the mean-squared displacement (MSD) of its nucleosomes over time and measure the nucleosome's diffusion coefficient $D$ which is given by

$$\mathrm{MSD}(t) = 6Dt \tag{1}$$

for particles that are freely diffusing in 3D space via Brownian motion. Shown in Fig. S4, $D = 8.22 \times 10^{-5}$ cm$^2$ / sim s and so $\tau = \frac{R_g^2}{D} \approx 2.55 \times 10^{-8}$ sim s, where sim s denotes simulation seconds.

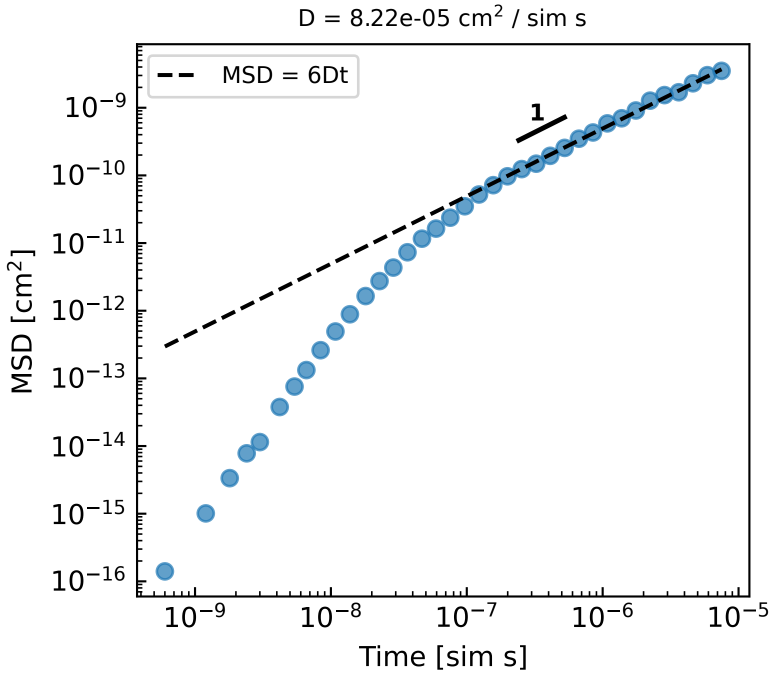


Figure S4: Mean-squared displacement (MSD) over time for a single 12-mer nucleosomal array with a linker DNA length of 25 bp. A slope of 1 is shown on the graph which indicates diffusive behavior.

Lastly we calculate $\tau$ for a physical system so that we can relate our simulated time to real, physical timescales. Because experiments have only measured the diffusion coefficients of nucleosomal arrays containing at most 4 nucleosomes with a linker length of 60 bp,[S3–S6] we extrapolate from these measurements the diffusion coefficient for a 12-mer nucleosomal array with linker length 25 bp. We do so by using the Einstein relation for dilute particles undergoing Brownian motion:[S7]

$$\xi = m\lambda_{real} = \frac{k_B T}{D} \tag{2}$$

where $\xi$ is the friction coefficient, $m$ is the mass of the particle, $T$ is temperature, $k_B$ is Boltzmann's constant, and $\lambda_{real}$ is the friction coefficient for the real system. We then assume that $\lambda_{real}$ is constant and estimate that a real 12-mer nucleosomal array with linker length 25 bp has a diffusion coefficient of $5.6 \times 10^{-8}$ cm$^2$/s. This indicates that a single second of simulation time is roughly equal to 1500 seconds in real time which means that $\tau$ is approximately 40 µs. We note that due to the assumptions inherent to this analysis, the precise timescales that we consider in our simulations should be considered approximate.

### Fitting 1CPN's nucleosome pair potential

1CPN can model specific post-translational modifications through the multiscale parametrization of its nucleosome pair potential (see Ref.[S1]). In the main text we utilize this feature to perform H4 tail acetylation and simulate chemically-detailed internucleosome interactions. In greater detail, 1CPN's nucleosome pair potential is parametrized by fitting a Zewdie potential to nucleosome pair potentials estimated by umbrella samplings of the near-atomistic 3SPN-AICG nucleosome model,[S8–S13] as done in the original 1CPN paper.[S1] Because the 3SPN-AICG model has residue resolution, 3SPN-AICG can be post-translationally modified directly which enables 1CPN to examine any post-translational modification at the large length scales of chromatin phase separation.

When we simulated the H4 tailless Zewdie potential from the original 1CPN paper,[S1] which Moller et al. showed to be energetically equivalent to H4 acetylation, we observed erroneous predictions for how H4 acetylation affected chromatin phase separation. We consequently investigated the fitting of the $\epsilon$ parameters in this H4 acetylated Zewdie potential and found the fit to be undetermined. To show this undetermined fitting, we first calculate a metric for a given Zewdie potential fit called the Zewdie sum $\Delta U_{\text{Zewdie}}$ which is given by the following equation:

$$\Delta U_{\text{Zewdie}} = \sum_{a \in A} \left( U_{\text{Zewdie,new}}(a) - U_{\text{Zewdie}}(a) \right) \tag{3}$$

where $A$ is the set of nucleosome-nucleosome orientations and distances from a single frame of a bulk condensate simulation, $U_{\text{Zewdie}}(a)$ is 1CPN's default Zewdie potential evaluated for a nucleosome-nucleosome interaction given by $a$, and $U_{\text{Zewdie,new}}(a)$ is the same but for the newly fit Zewdie potential. Effectively, the Zewdie sum estimates how the total system energy would change for a new parametrization. We find that Zewdie potential fits with varying initial conditions can result in different values of the Zewdie sum and therefore different values of the overall angle-dependent pair potential (Fig. S5).

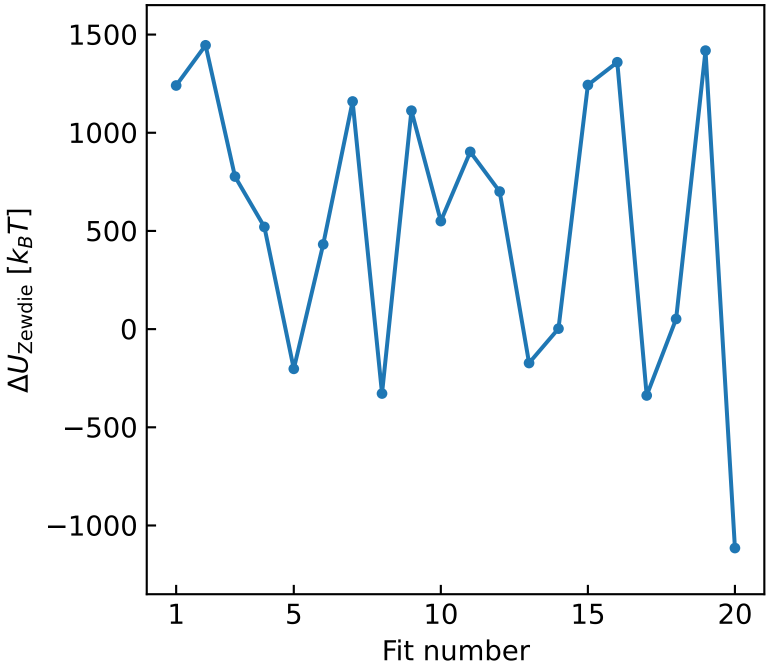


Figure S5: The Zewdie sums for Zewdie potential fits to the H4 acetylated 3SPN-AICG nucleosome pair potentials. The pair potentials used in the fitting correspond to face-face, face-side, and side-side interaction types. Each fit has free parameters $\epsilon_{000}$, $\epsilon_{cc2}$, $\epsilon_{220}$, and $\epsilon_{222}$, and uses random initial conditions.

In the main text, we used the unmodified Zewdie potential fit from the original 1CPN paper[S1] and a non-unique H4 acetylated Zewdie potential fit (see Table S1). After later informal discussions with Yiheng Wu and Aria Coraor (U Chicago), we found that by keeping $\epsilon_0$ constant as well as including the rotated face-side orientation, the H4 acetylated Zewdie potential fit became stable (i.e. unique) under different initial conditions. This is shown in Fig. S6. Reassuringly, we find that the parameter values and Zewdie sums of this unique fit and our non-unique fit are very close in value (Table S1), which reveals that they predict similar equilibrium properties.

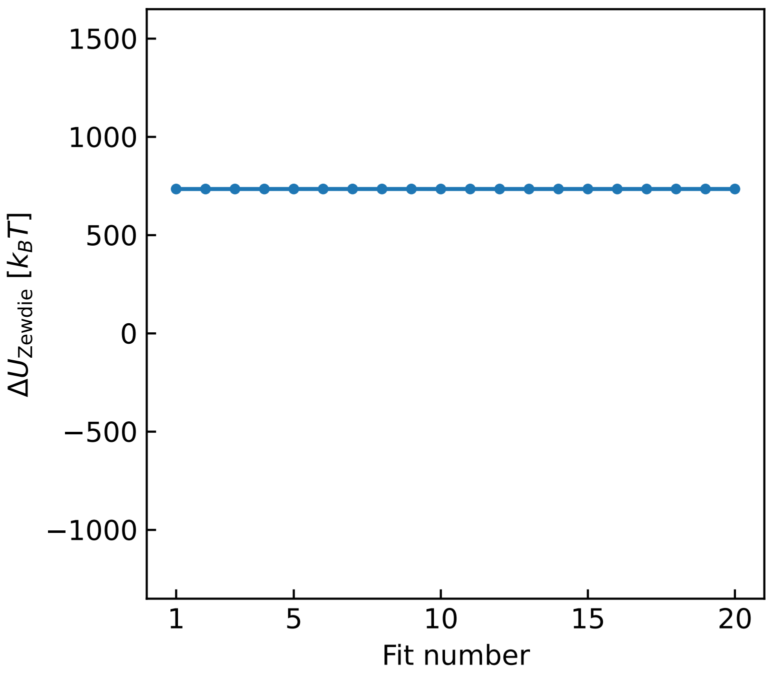


Figure S6: The Zewdie sums for Zewdie potential fits to the H4 acetylated 3SPN-AICG nucleosome pair potentials. The pair potentials used in the fitting correspond to face-face, face-side, side-side, and rotated face-side interaction types. Each fit has free parameters $\epsilon_{000}$, $\epsilon_{cc2}$, $\epsilon_{220}$, and $\epsilon_{222}$, and uses random initial conditions.

**Table S1: H4 acetylated Zewdie parameters**

| | Non-unique H4 acetylated fit [a] | Unique H4 acetylated fit |
|---|---|---|
| $\sigma_0$ | 55.00 | 55.00 |
| $\sigma_{000}$ | 1.564 | 1.564 |
| $\sigma_{cc2}$ | -0.7538 | -0.7538 |
| $\sigma_{220}$ | 0.1512 | 0.1512 |
| $\sigma_{222}$ | 0.2670 | 0.2670 |
| $\sigma_{224}$ | 0 | 0 |
| $\epsilon_0$ | 1.164 | 1.164 |
| $\epsilon_{000}$ | 0.4767 | 0.5264 |
| $\epsilon_{cc2}$ | 0.4996 | 0.4003 |
| $\epsilon_{220}$ | -0.06983 | -0.1773 |
| $\epsilon_{222}$ | -0.7918 | -0.4168 |
| $\epsilon_{224}$ | 0 | 0 |
| Zewdie sum | 1268 | 734.8 |

[a] Used in our work to model H4 acetylation.

### Mapping nucleosome sites to whole nucleosomes

1CPN represents nucleosomes using a single anisotropic site that reproduces the excluded volume of a nucleosome through the Zewdie potential. This nucleosome site is located at the center of mass of each nucleosome, which allows us to conveniently calculate the mean-squared displacement of nucleosomes and the nucleosome-nucleosome structure factor. However, when quantifying the dimensions of a nucleosomal array (as done in the main text when we approximate them as cylinders) we found that a slightly more accurate representation of the nucleosome's excluded volume was necessary. In our approach, we place an additional number of ghost sites randomly inside of the nucleosome's excluded volume. Shown in Fig. S7, we see that adding 100 ghost sites well-approximates the nucleosome's excluded volume. We note that these ghost sites were only added as a post-processing step when analyzing 1CPN trajectories and that these ghost sites were not present in the original 1CPN molecular dynamics simulations.

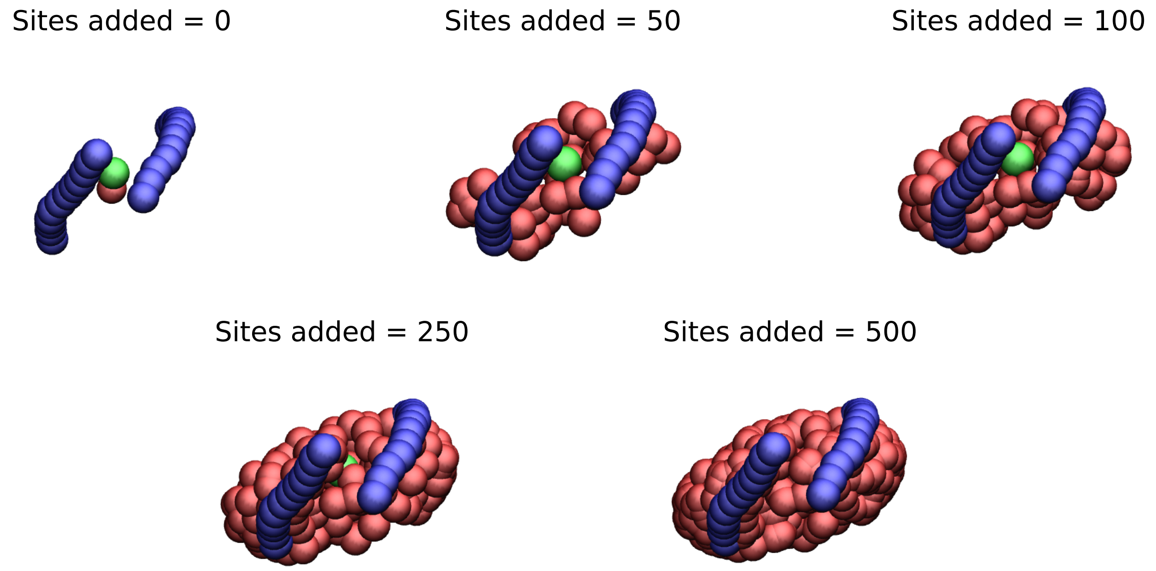


Figure S7: Site-based VMD[S14] visualization of 1CPN nucleosomes with different numbers of additional ghost sites. A sites added equal to 0 corresponds to a visualization containing just the original nucleosome site.

(S7) Allen, M. P.; Tildesley, D. J. *Computer simulation of liquids*; Oxford university press, 2017.

(S8) Li, W.; Wolynes, P. G.; Takada, S. Frustration, specific sequence dependence, and nonlinearity in large-amplitude fluctuations of allosteric proteins. *Proceedings of the National Academy of Sciences* **2011**, *108*, 3504–3509.

(S9) Freeman, G. S.; Lequieu, J. P.; Hinckley, D. M.; Whitmer, J. K.; De Pablo, J. J. DNA shape dominates sequence affinity in nucleosome formation. *Physical Review Letters* **2014**, *113*, 168101.

(S10) Freeman, G. S.; Hinckley, D. M.; Lequieu, J. P.; Whitmer, J. K.; De Pablo, J. J. Coarse-grained modeling of DNA curvature. *The Journal of chemical physics* **2014**, *141*.

(S11) Lequieu, J.; Córdoba, A.; Schwartz, D. C.; de Pablo, J. J. Tension-dependent free energies of nucleosome unwrapping. *ACS central science* **2016**, *2*, 660–666.

(S12) Lequieu, J.; Schwartz, D. C.; de Pablo, J. J. In silico evidence for sequence-dependent nucleosome sliding. *Proceedings of the National Academy of Sciences* **2017**, *114*, E9197–E9205.

(S13) Moller, J.; Lequieu, J.; de Pablo, J. J. The free energy landscape of internucleosome interactions and its relation to chromatin fiber structure. *ACS central science* **2019**, *5*, 341–348.

(S14) Humphrey, W.; Dalke, A.; Schulten, K. VMD: visual molecular dynamics. *Journal of molecular graphics* **1996**, *14*, 33–38.
